# Evolution of recombination landscapes in diverging populations of bread wheat

**DOI:** 10.1101/2021.01.22.427740

**Authors:** Alice Danguy des Déserts, Sophie Bouchet, Pierre Sourdille, Bertrand Servin

**Author notes:** Corresponding authors Pierre Sourdille;, Bertrand Servin.

## Abstract

Reciprocal exchanges of DNA (crossovers) that occur during meiosis are mandatory to ensure the production of fertile gametes in sexually reproducing species. They also contribute to shuffle parental alleles into new combinations thereby fuelling genetic variation and evolution. However, due to biological constraints, the recombination landscape is highly heterogenous along the genome which limits the range of allelic combinations and the adaptability of populations. An approach to better understand the constraints on the recombination process is to study how it evolved in the past. In this work we tackled this question by constructing recombination profiles in four diverging bread wheat (*Triticum aestivum L*.) populations established from 371 landraces genotyped at 200,062 SNPs. We used linkage disequilibrium (LD) patterns to estimate in each population the past distribution of recombination along the genome and characterize its fine-scale heterogeneity. At the megabase scale, recombination rates derived from LD patterns were consistent with family-based estimates obtained from a population of 406 recombinant inbred lines. Among the four populations, recombination landscapes were significantly positively correlated between each other and shared a statistically significant proportion of highly recombinant intervals. However, this comparison also highlighted that the similarity in recombination landscapes between populations was significantly decreasing with their genetic differentiation in most regions of the genome. This observation was found to be robust to SNP ascertainment and demography and suggests a relatively rapid evolution of factors determining the fine-scale localization of recombination in bread wheat.

## Introduction

Meiotic recombination (or crossover; CO) is the obligate genetic exchange between homologous chromosomes that occurs during the production of gametes in sexually reproducing species. Besides its role in ensuring proper segregation of chromosomes in gametes, it also impacts evolution by breaking linkage between advantageous and deleterious alleles and by creating novel combinations of alleles (Barton 1995; Charlesworth and Barton 1996; Otto 2009). Recombination rates are highly variable between species and also at different genomic scales. At the chromosomal level, COs are not evenly distributed depending on either the size of the chromosomes, the region of the chromosomes or on interference. Interference was first observed in Drosophila (for review see (Berchowitz and Copenhaver 2010) and is defined as the impossibility for a type I CO (*i*.*e*. COs that are submitted to interference contrary to type II COs that are not) to occur in the vicinity of another CO from the same type. Type I COs are thus more regularly spaced along chromosomes than expected from random (Zickler and Kleckner 2015). Within chromosomes, some regions are also deprived of Cos, such as centromeres where COs are absent in all species studied so far. Moreover, in many species, distribution of COs is skewed towards telomeres where they show a high tendency to occur. In wheat (*Triticum aestivum* L.) for example, more than 80% of the recombination events occur in the terminal regions of the chromosomes representing less than 20% of the genome (Saintenac et al. 2009; Choulet et al. 2014; Darrier et al. 2017; International Wheat Genome Sequencing Consortium IWGSC 2018). The main hypothesis to explain this behaviour is the early initiation of synapsis and recombination in the telomeric regions as shown in barley (Higgins et al. 2012; Dreissig et al. 2019). In species with small chromosomes such as *Arabidopsis thaliana* or rice (*Oryza sativa*), recombination events are more evenly distributed along the chromosomes with the exception of the centromeres (Choi et al. 2013; Drouaud et al. 2013; Marand et al. 2019). However and despite these differences, there is rarely more than three COs per chromosome and per meiosis in every species (Mercier et al. 2015).

At a local scale, in most species including yeast, birds, snakes, fishes, mammals and plants, COs mainly occur in small regions of a few kilobases (kb) called hotspots (Myers et al. 2005; Mancera et al. 2008; Choi and Henderson 2015; Singhal et al. 2015; Shanfelter et al. 2019; Schield et al. 2020). In some mammals, these hotspots are determined by PRDM9, a SET-domain protein with a zinc-finger array that binds DNA (Boulton et al. 1997; Oliver et al. 2009; Baudat et al. 2010; Myers et al. 2010). PRDM9 recognizes specific DNA motifs and deposits an epigenetic landmark (histone H3 trimethylated on lysine 4: H3K4me3) that is further recognized by the machinery forming double-strand breaks that initiates COs (Murakami et al. 2020). However, many if not most species (*e*.*g*. birds, plants, yeast, snakes and fishes) do not exhibit a PRDM9 derived mechanism for driving the localization of recombination hotspots. There are mainly determined by chromatin features and are often found in accessible chromatin regions (Auton et al. 2013; Choi and Henderson 2015; Singhal et al. 2015; Marand et al. 2017; Marand et al. 2019) although intermediate situations exist (Schield et al. 2020).

Determinisms of local recombination rates with regards to the distribution of CO hotspots remain unknown in many organisms. One approach to better understand these determinisms is to characterize the evolution of the recombination landscape and evidence its conservation or lack thereof. This can be achieved by contrasting recombination landscapes in closely related species (Stapley et al. 2017) or in differentiated populations of the same species (Kong et al. 2010; Salomé et al. 2012; Petit et al. 2017). For example, in rice, less than 20% of the CO hotspots are common between the two subspecies *Oryza sativa ssp. japonica* and *O. s. ssp. indica* (Marand et al. 2019) although they diverged relatively recently (440,000-86,000 years ago (YA); Ma and Bennetzen 2004; Vitte et al. 2004; Zhu and Ge 2005; Tang et al. 2006). Similarly, analysis of the recombination landscapes in the cocoa-tree (*Theobroma cacao*) showed only little overlap of recombination hotspots across ten diverging populations with less divergent populations showing higher level of overlap (Schwarzkopf et al. 2020). An analysis of the recombination landscapes in wild barley (*Hordeum vulgare ssp. spontaneum*) vs. domesticated barley (*H. vulgare*) revealed that recombination tend to cluster in more distal regions in the latter (Dreissig et al. 2019) while the two species diverged approximately 4 million years ago (MYA; Brassac and Blattner 2015), and domestication began approximately 10,000 YA (Badr et al. 2000). A finer-scale analysis among subpopulations of wild barley revealed that recombination rate varied according to environmental conditions (temperature, aridity, solar radiation, annual precipitations), suggesting that environmental factors might explain part of these differences (Dreissig et al. 2019).

High-density genotyping SNP arrays as well as new generation sequencing (NGS) approaches now allow to analyse large collections of wild/domesticated, ancient/modern populations of both animals and plants. Such a large amount of accurate data permits to better decipher the recombination landscape from patterns of Linkage Disequilibrium (LD) (Li and Stephens 2003; Auton and McVean 2007; Chan et al. 2012). The advantages of using this approach stem from the large number of meiosis that occurred during the evolution of sampled populations compared to bi-parental or multi-parental experimental populations. First, as LD-based recombination inference is based on recombination happening in many different individuals it should consequently be less sensitive to individual specific variation, which might occur in the presence of structural variation for example (Bauer et al. 2013; Rowan et al. 2019). Second, LD-based recombination rate estimates are more resolutive as genetic diversity is higher compared to experimental segregating populations that typically involve few parents. However, the drawback of this approach is that the recombination landscapes obtained can be affected by evolutionary forces (Charlesworth and Charlesworth 2010; Auton and McVean 2012; Choi and Henderson 2015) and consequently have to be interpreted cautiously.

Despite these limitations, the LD-based approach was successfully applied at the whole-genome level in many species including birds (Singhal et al. 2015; Smeds et al. 2016), yeast (Tsai et al. 2010), Arabidopsis (Choi et al. 2013), rice (Marand et al. 2019), barley (Dreissig et al. 2019) and bread wheat chromosome 3B (Darrier et al. 2017). This latter study was limited to chromosome 3B as it was the only chromosome presenting a sufficiently high-standard reference sequence at that time (Choulet et al. 2014; IWGSC 2014). The analysis of two collections representative of the Asian and European genetic pools revealed a high similarity between their recombination profiles. These LD-based profile were also shown to be consistent with a meiotic recombination profile derived from a bi-parental population (Chinese Spring x Renan; Choulet et al. 2014). This result suggested that recombination rate estimation through a LD-based approach could be even more informative and resolutive along the whole genome using the last gold-standard reference sequence available (IWGSC 2018), as well as high-density genotyping of large wheat collections.

The complexity and huge size (16 gigabases) of the wheat genome have long hampered the development of high throughput genomic tools as well as the establishment of a whole genome sequence. Bread wheat is an allo-hexaploid species (AABBDD; 2n = 6x = 42) derived from two successive interspecific crosses involving three diploid species (for details, see https://www.wheatgenome.org/; IWGSC 2014; IWGSC 2018): *T. monococcum ssp. urartu* (AA genome), a yet-unknown species related to the *Sitopsis* section (SS genome related to the wheat BB genome) and *Aegilops tauschii* (DD genome). However, international efforts combined with appropriate and original strategies using chromosome sorting, chromosome-specific BAC libraries, paired-end short-read sequencing and relevant assembly approaches, lead to the publication of a high-standard, annotated, oriented and anchored sequence of the wheat genome (IWGSC 2018). At the same time and despite the presence of a high proportion of transposable elements (85%; Wicker et al. 2018), high-density SNP arrays have been successfully developed and used for marker-assisted selection (Sun et al. 2020) and for the characterization of collections (Winfield et al. 2016; Balfourier et al. 2019). In the study of Balfourier et al. (2019), the genetic structuration of 4,506 bread wheat landraces and cultivars representative of the worldwide diversity was described using the TaBW 280k SNP chip. These data offer the opportunity to extend previous work on bread wheat by analysing recombination along the whole genome and across more populations. We compared the ancestral recombination profiles of four populations with the meiotic recombination observed in a biparental recombinant inbred lines (RILs) population (Chinese Spring x Renan; CsRe). We developed specific statistical models to evaluate and minimize the influence of evolutionary forces on the comparison of recombination landscapes between populations.

## Results

### Identification of four populations of bread wheat landraces

Establishing LD-based recombination maps requires samples of unrelated chromosomes from a homogeneous population. We extracted a subset of 371 landraces representative of the worldwide diversity from Balfourier et al. (2019), forming four distinct and mostly homogeneous genetic populations (see methods) (Figure 1) that were named according to the geographical origins of their members: the West-European population (WE), composed of 127 accessions originating from France (52 accessions), Spain (10), Germany (8) and from 30 other Western European, Mediterranean countries and Iberian peninsula; the East-European population (EE), composed of 70 accessions originating from France (9), the Russian Federation (7), Ukraine (5) and from 27 other Eastern European countries; the West-Asian population (WA), composed of 97 accessions originating from Afghanistan (8), Pakistan (8), Turkey (8) and from 33 other of Caucasian and Central Asia countries and Indian peninsula; the East-Asian population (EA) composed of 77 accessions originating from China (61), Japan (7), the Republic of Korea (4) and from 5 other South East Asian countries (supplementary file S1). The genetic differentiation of the four populations confirmed an increasing genetic divergence along a Eurasian gradient (Figure 1), consistent with isolation by distance, selection and differentiation that occurred during the initial independent spreads of bread wheat from the Cradle of Agriculture and Wheat in the Fertile Crescent toward Europe on the one hand and Asia on the other hand during the Neolithic period (Balfourier et al. 2019). WE and EE are the most related groups (F_ST_ = 0.015) while WE and EA are the more divergent ones (F_ST_ = 0.085) and also the most geographically distant. The WA population is the closest population to the tree root possibly because it includes accessions that were collected not far from the centre of domestication of bread Wheat (Fertile Crescent: Turkey, Iraq, Iran; Caucasus and Caspian Sea: Armenia, Georgia, Kazakhstan, Turkmenistan). The EA population appears as a very differentiated and homogenous population. WE and EE are less differentiated because they separated more recently from each other (Balfourier et al. 2019).

**Figure 1:**
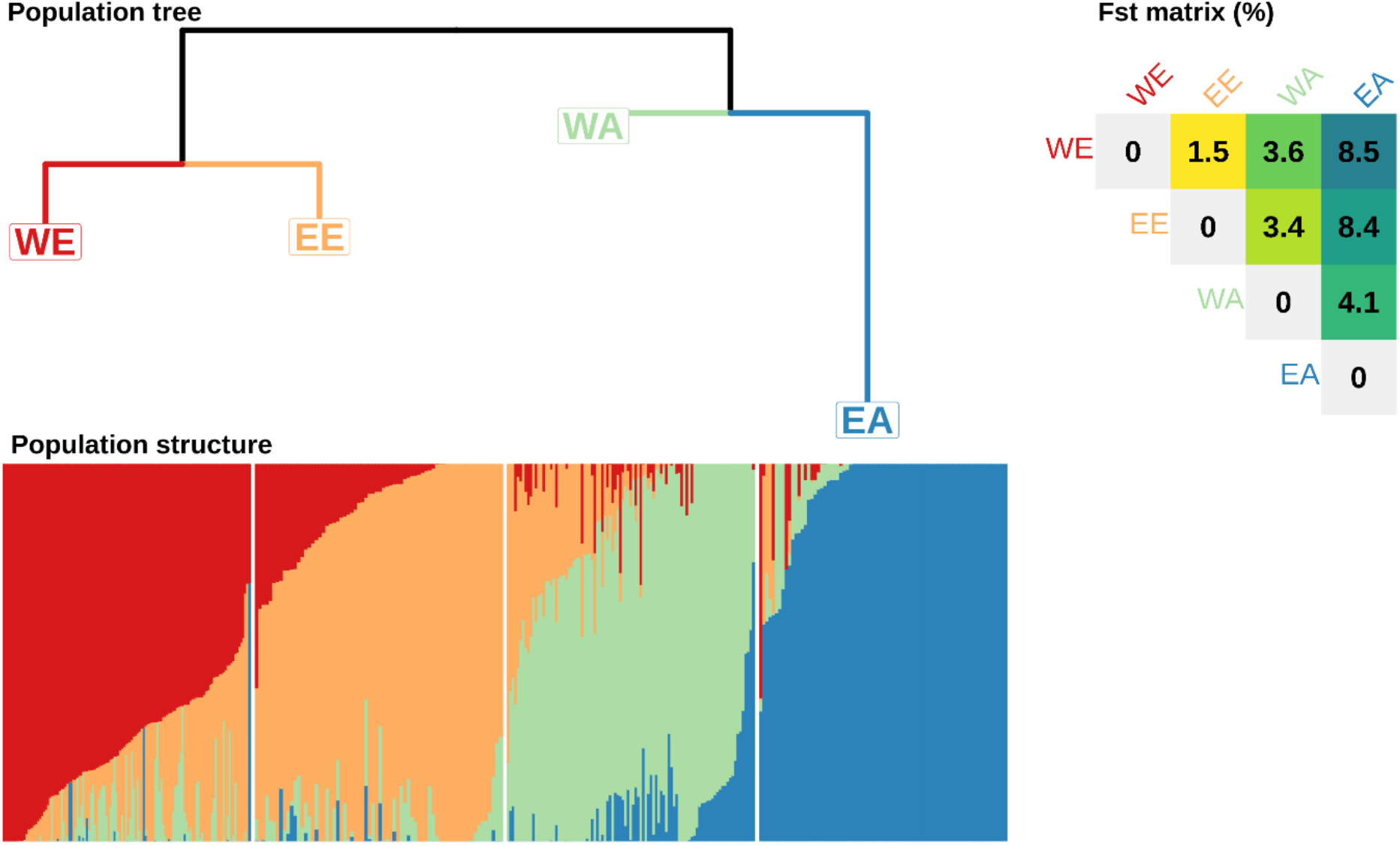
Bread wheat landrace genetic divergence and structuration. **Population tree** Neighbour Joining tree built with pairwise Reynold distance matrix computed on SNP alleles and rooted by HAPFLK software (Bonhomme et al. 2010; Fariello et al. 2013). WE = West Europe, EE = East European, WA = West Asia, EA = East Asia. **Fst matrix (%)** Weir and Cockerham pairwise Fst computed with simple matching distance of haplotypic alleles. **Population structure** Admixture coefficients for K=4 from Balfourier et al. (2019) using STRUCTURE software and haplotypic alleles.

The genetic composition of the four populations appeared quite distinct between populations but homogenous within populations when described by the K=4 admixture analysis of Balfourier et al. (2019) (Figure 1). WE, EE, WA and EA have almost all their members belonging to the same specific dominant group (respectively named by Balfourier et al. (2019) as North West European, South East European, Central Asian and African and South East Asian groups) with a high membership coefficient: 0.74 on average for WE (standard deviation = 0.16), 0.81 for EE (±0.16), 0.73 for WA (±0.17) and 0.93 for EA (±0.14). The WE and WA populations appear to be more admixed than EE and EA at K=8 (supplementary Figure S1). In order to analyse groups that are large enough to estimate relevant statistics, we split landraces into four populations, although there is some sub-structuration within populations. This was motivated by the fact that model we used to estimate LD-based recombination rates was shown to be robust to moderate levels of structuration (Li and Stephens 2003).

### Recombination patterns are broadly conserved across populations

#### Robust meiotic recombination map of a population of Recombinant Inbred Lines (RILs)

We established a meiotic recombination map from recombination events observed in a population of 406 F6 RILs (termed CsRe in the following). This population is derived from a cross between two bread wheat varieties: Chinese Spring and Renan belonging respectively to the EA and WE gene pools. The CsRe population was previously genotyped for the same set of SNPs as the landraces (Rimbert et al. 2018). Recombination rates in CsRe were derived from the observed proportion of recombinants in each of the 79,543 intervals defined by SNPs that were polymorphic in the cross. The distribution of recombinants in these intervals led to extremely contrasted situations. On one hand, 60% of these intervals harboured no recombinant among the 406 offspring. On the other hand, a few recombinants were observed in very small intervals. Using a simple statistical approach to estimate recombination rates from these observation produces extreme differences in recombination rates that are highly influenced by the limited sample size available. In order to produce more reliable estimates that better account for sample size and uncertainty, we fitted a Bayesian Poisson Gamma model on the observed recombinant counts (see methods). With this model, the estimates of recombination rates in the RILs population ranged from almost 0 to 78 cM/Mb among intervals. Compared to the simple estimates that ranged up to 2,806 cM/Mb this approach has the advantage of shrinking extreme values that are unrealistic and solely due to the limited number of RILs available. Consistent with the Bayesian model correcting for the effect of sample size, the correlation between naive and Bayesian estimates increases with the number of observed recombinants per intervals (Supplementary Figure S2), *i*.*e*. the two approaches converge to the same inference when the data is informative enough.

#### Validation of LD-based recombination maps on CsRe meiotic recombination map

LD-based recombination maps were inferred from patterns of LD between polymorphic SNPs for each landrace population independently using PHASE (Li and Stephens 2003). As LD is strongly related to meiotic recombination but can also result from evolutionary forces, those maps were compared to the meiotic CsRe recombination map described above.

Before estimating LD-based recombination rates, SNPs were filtered out on Minor Allele Frequency (MAF) with a minimum value of 3% within each population, yielding to 170,509 SNPs for WE, 161,137 for EE, 171,901 for WA and 131,585 for EA. The average marker density was 11 SNPs/Mb with most of the SNPs located at telomeres (25 SNPs/Mb) while centromeres were depleted in SNPs (3 SNPs/Mb). SNP density was almost three times higher on the A and B genomes compared to the D genome (respectively 14, 14 and 5 SNPs/Mb). This is consistent with the lower rate of polymorphism of the wheat D genome.

Both LD-based and meiotic recombination profiles showed the same global patterns at the chromosome scale (Figure 2). In both approaches, the telomeric regions R1 and R3 of chromosomes showed recombination rates (average LD-based recombination rate in WE = 1e-2/kb; average CsRe Bayesian recombination rate = 0.8 cM/Mb) around ten times higher than the pericentromeric regions R2a and R2b (2e-3/kb; 0.1cM/Mb) and one hundred times higher than the centromeric regions C (2e-4/kb; 0.01 cM/Mb). Recombination rates on the D genome were higher than recombination rates in the A and B genomes. This can be explained by a lower genetic diversity facilitating homologous pairing and recombination during meiosis (Saintenac et al. 2011; Rimbert et al. 2018) and by the reduced physical size of D chromosomes which, associated to an obligatory crossing over per tetrad, results in higher recombination rates on smaller chromosomes.

**Figure 2:**
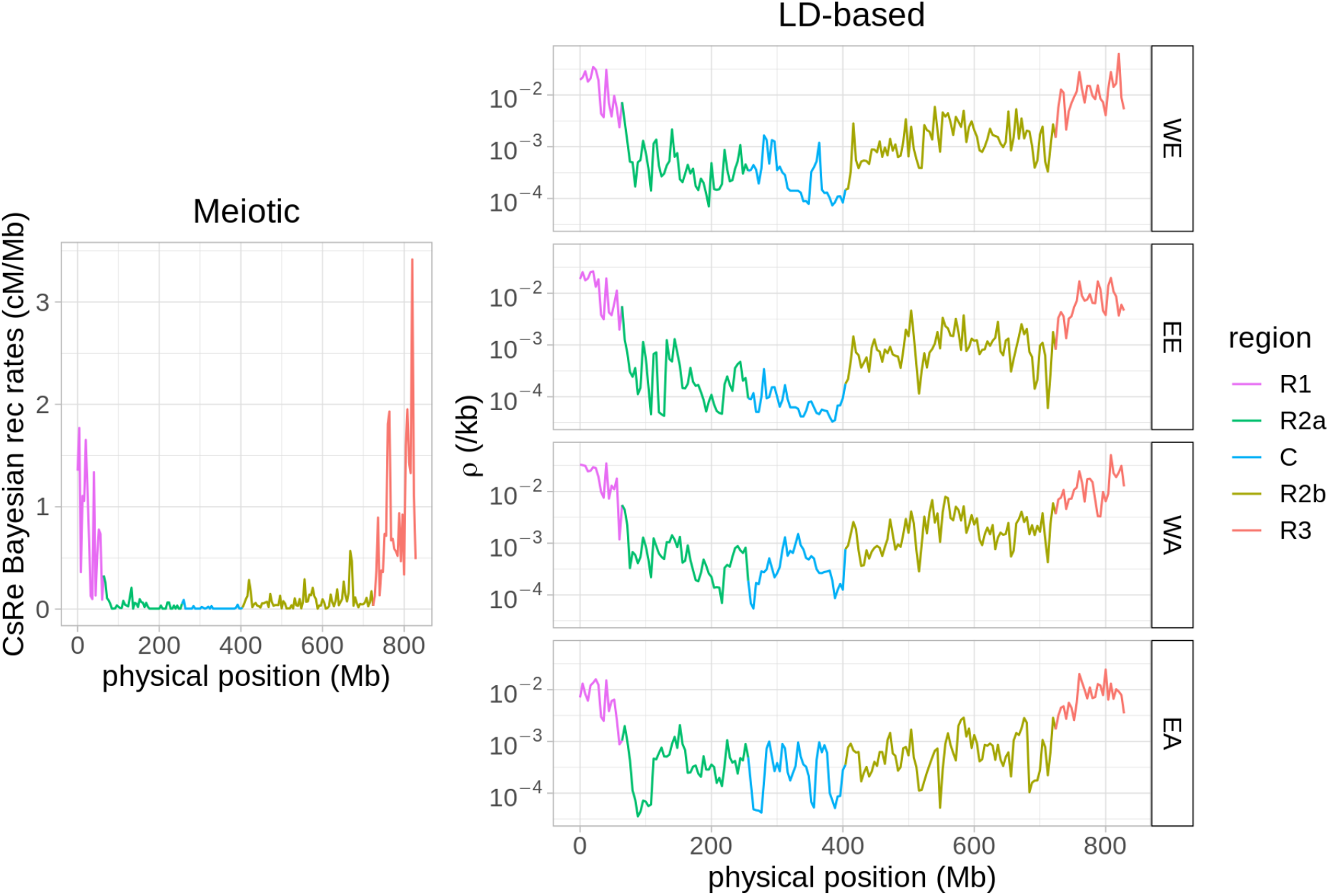
Meiotic and LD-based recombination profiles in 4 Mb windows along chromosome 3B in the CsRe segregating population (left) and in the four West European (WE), East European (EE), West Asia (WA) and East Asia (EA) collections (right). Each colour corresponds to genomic regions defined by Choulet et al. (2014): highly recombining telomeres R1 (magenta) & R3 (red); low recombining pericentromeres R2a (dark green) & R2b (light green); and centromere C (blue) where recombination rates are close to 0. LD-based recombination profiles at natural scale are present in supplementary Figure S3.

The genome-wide correlation of LD-based recombination profiles and CsRe Bayesian meiotic recombination profile was quite high for the four populations (≥ 0.7, Table 1) but slightly higher for European populations (pairwise significant differences according Zou’s test (Zou 2007), R cocor package). These high correlations are explained by the strong partitioning of the recombination profile along chromosomes present in all bread wheat populations, *i*.*e*. low recombination rates in centromeres and high recombination rates in telomeres (Figure 3). Beyond this inter-region contrast, the within region correlations were lower but still significantly positive (1AR1 – 7DR3, Figure 4, supplementary file S2). In telomeres R1 & R3 and pericentromeres R2a & R2b, the average correlation ranged between 0.50 in EA to 0.58 in WE (Table 1), with an average of 0.56 all populations confounded.

**Table 1.** Correlation of the LD-based recombination profiles of the 4 populations of landraces with CsRe Bayesian meiotic recombination profile. Recombination rates were averaged in 4 Mb windows.

|  | WE | EE | WA | EA |
| --- | --- | --- | --- | --- |
| Genome-wide | 0.76 | 0.75 | 0.74 | 0.70 |
| Average on 84 genomic regions (R1, R2a, R2b, R3 of chr 1A – 7D) | $0.58 \pm 0.22$ | $0.55 \pm 0.28$ | $0.55 \pm 0.27$ | $0.50 \pm 0.29$ |
| Average on 21 C regions (chr 1A – 7D) | $0.32 \pm 0.33$ | $0.30 \pm 0.34$ | $0.20 \pm 0.34$ | $0.19 \pm 0.36$ |

**Figure 3:**
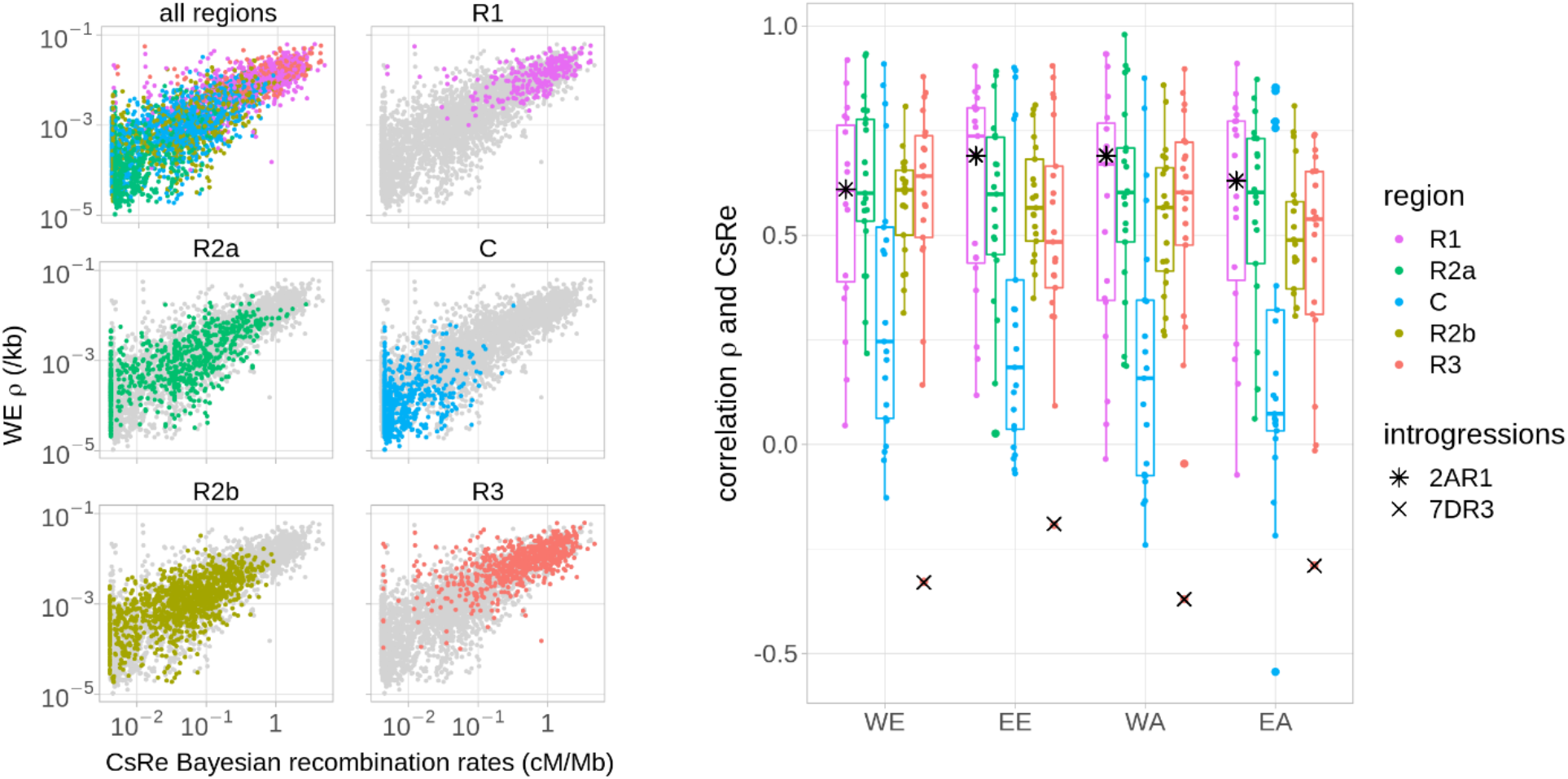
**Left** Genome-wide relationship between the CsRe biparental population meiotic recombination rates and the LD-based recombination profile of a Western European (WE) bread wheat population. Points represent the recombination rates averaged within 4 Mb windows. **Right** Correlation of LD-based and CsRe recombination rates at a 4 Mb scale within each genomic region of bread wheat genome (1AR1…7DR3). Genomic regions smaller than 20 Mb are not included. Stars represent genomic regions including well documented introgressions in CsRe population.

**Figure 4:**
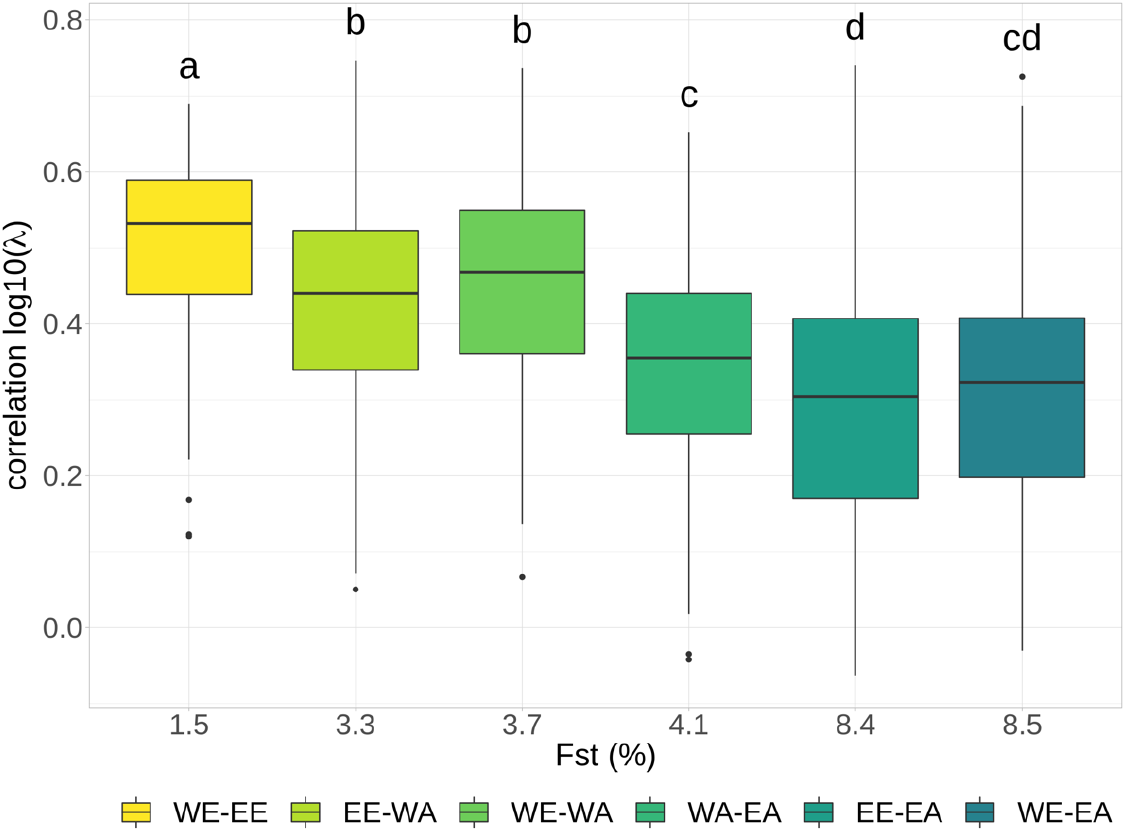
Relationship between pairwise correlation of LD-based recombination intensity λ and F_st_. Each boxplot contains 84 correlation coefficients corresponding to the 84 genomic regions (1AR1…7DR3, excluding centromeres). Letters above boxplots indicate if the means are significantly different between populations (Fisher test, p-value < 2.2e-16).

The recombination rates in centromeric regions showed much lower consistency: the correlation of centromeric LD-based recombination rates and CsRe recombination rates ranged from 0.19 in EA to 0.32 in WE. Considering the low correlation but also the low SNP density and the fact that centromere sequence assemblies are challenging because of the presence of numerous repeated sequences such as transposons and retro-transposons (IWGSC 2018; Wicker et al. 2018), centromeres were no longer included in the analyses.

Among the genomic regions considered, the 7DR3 one exhibited a strikingly low and negative correlation between LD-based and meiotic recombination rates in all populations (≤ −0.19, Figure 4). This result is due to a low recombination rate in part of this region in the CsRe biparental genetic map that is not observed in LD-based maps (supplementary Figure S4). This low recombination rate can be explained by the fact that the Renan line carries an inter-specific introgression of 28 Mb on chromosome 7D around the eyespot resistance gene *Pch1* coming from *Aegilops ventricosa* (tetraploid species; DDNN) (Maia 1967). This introgression does not recombine in the CsRe cross as this was previously evidenced in another background (Worland et al. 1988). Interestingly the Renan line carries another 20 Mb introgression from *Aegilops ventricosa* in 2AR1 region around the *Lr37/Sr38/Yr17* resistance gene cluster. However, in this region, contrary to the 7DR3 case, the LD patterns are also consistent with an overall low recombination rate. Since the introgression in region 2AR1 suppresses recombination in an already low recombining region, this explains why the correlation coefficient with LD-based profiles in 2AR1 does not stand out particularly (Figure 4).

Both CsRe and LD-based maps show a high heterogeneity in the distribution of recombination rates along chromosomes: on average 36% of physical distance represents 80% of genetic distance in all our populations. To further study the distribution of chromosome sites cumulating historical crossovers, we defined highly-recombining intervals (HRIs) in the four landrace populations as intervals with an LD-based recombination rate exceeding four-times the background recombination rate (λ ≥ 4, see Methods). Combining all four populations, this resulted in 8,713 HRIs, with a median deviation to background recombination rate λ = 6.5 (range λ = 4 to λ = 511). Note that we avoid here the term LD-based recombination *hotspot* as functional hotspots typically span much smaller genomic regions (size < 5 kb, (Marand et al. 2019)) than our defined HRIs (median size = 20 kb). Therefore, we cannot be sure that an HRI harbours a single recombination hotspot. The repartition of HRIs along the genome was heterogeneous. Most HRIs (73%) were located in telomeric R1 or R3 regions, and the other HRIs (27%) in pericentromeric R2a or R2b regions. As HRIs corresponded to respectively 2% and 1% of intervals in those regions, telomeres were significantly enriched in HRIs compared to pericentromeres (significant chisq test, P-value < 2.2e-16). These HRIs represented 15 % of LD-based genetic distance (from 12 % in EA to 18% in WA) and around 9% of the physical distance (from 6% in EA to 10% in WE). The proportion of HRIs and non-HRIs intervals co-localizing with open-chromatin features such as genes, 5’UTR and 3’UTR features was measured independently within regions R1, R2a, R2b and R3 and then averaged. It revealed that HRIs are much more co-localizing with gene features than non-HRIs intervals, as the proportion of HRIs co-localizing with genes, 5’UTR and 3’UTR was respectively 80%, 56% and 54%, but dramatically decreased to 53%, 22% and 24% when considering non-HRIs intervals (supplementary Figure S5). Consequently, 80% of HRIs were associated with gene features versus 53 % of the other intervals. The density of HRIs is also positively associated with the CsRe meiotic recombination rate averaged in 4 Mb windows in each genomic region R1, R2a, R2b and R3 (P-value < 2.2e-16). The proportion of CsRe crossovers overlapping HRIs ranged from 20% in EA to 37% in WE. Most HRIs (82%) overlapped at least one CsRe crossover.

Despite high similarities between LD-based and meiotic recombination profile within genomic region, there is still the possibility that LD-based recombination rates might be locally influenced by evolutionary forces, such as positive selection, as shown by (Petit et al. 2017) in sheep for example. To evaluate the potential effects of positive selection on the LD-based maps, we studied whether a set of genes known to be involved in domestication (*e*.*g*. brittle rachis (*Brt*), tenacious glume (*Tg*), homoeologous pairing (*Ph*) or non-free-threshing character (*Q*)) or recent crop improvement (Pont et al. 2019) were found in regions outliers for the ρ/CsRe ratio. The results showed no evidence of reduced recombination around these genes (supplementary Figure S6).

As no strong bias of evolutionary forces was evidenced, we converted meiotic recombination map specific to each landrace population in supplementary file S3. Briefly, LD-based recombination rates being proportional to landraces specific meiotic recombination rates, we removed the proportionality coefficient by rescaling those LD-based map on CsRe Bayesian recombination map (supplementary protocol S1). Generally, the LD-based profiles of recombination are congruent with the meiotic recombination map in the CsRe RILs population. This validates the use of LD-based recombination maps to study the evolution of recombination patterns.

### Significant differences between LD-based population-specific recombination maps

Our results reveal that the average LD-based recombination rates vary in a two-fold range between populations: WE has the highest rate and EA the lowest (WE: ρ = 0,004/kb; WA: ρ = 0,004/kb; EE ρ = 0,003/kb; EA: ρ = 0,002/kb; excluding centromeres). This ranking between populations could be explained by genetic diversity levels (Figure 1) as well as by different average meiotic recombination rates. The fact that WE and WA are more admixed populations than EE and EA favoured a more important contribution of diversity levels compared to a real difference on average recombination intensity. To eliminate the systematic effect of diversity and demography on recombination rate estimates, we chose to compare the population recombination profiles in terms of the deviation from their local background recombination rates. Specifically, the Li and Stephens’s model (2003) estimates an interval specific recombination parameter (λ) that measures the relative rate of recombination of an interval compared to its neighbours in a 2 cM window (see Methods). We therefore expect population-specific effects (other than local variation in recombination) to affect the background recombination rate but not the relative intensities of intervals measured by the parameter λ.

The similarity of λ profiles along the genome was evaluated by fitting a linear mixed model on the variations of 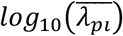 within each genomic region. In almost all genomic regions (79 out of 84), a linear model specifying a variance-covariance matrix with different correlation coefficients for each pair of populations showed a lowest BIC than a simpler model specifying a variance-covariance matrix including only one common correlation coefficient for all pairs of populations (see Methods). This indicates that local variations of recombination rates are significantly different between populations.

The average correlation of local variations of recombination rates across genomic regions was twice higher for the highest correlated pair WE-EE (0.47 ± 0.11) than for the lowest one EE-EA (0.20 ± 0.11), with an average value of 0.32 (Figure 4). Such decrease in the genome-wide similarity of recombination rates can also be measured using a Gini coefficient (Gini 1936). The Gini coefficient is a measure of the unevenness of a distribution. It is best known for its use in economics to measure the repartition of wealth among individuals. A Gini coefficient of 0 corresponds to a uniform distribution and one of 1 corresponds to the case where the distribution is a single point mass. Here we use the Gini coefficient to measure the unevenness of the repartition of genetic distance along chromosomes between two different populations by computing it based on the distribution of recombination in one population along the genetic map of the other: a Gini coefficient of 0 corresponds to identical recombination profiles and the more divergent the distribution in recombination profiles is, the higher Gini coefficient is. In our case, the pairwise Gini coefficients increased along the Eurasian gradient, with lower values for closely related population (around 0.43 for WE-EE) and higher values in distant populations (0.77 for WE-EA), meaning that similarity in distribution of LD-based genetic distance along chromosomes decreases along the Eurasian gradient (supplementary Figure S7).

In light of these significant differences in the local repartition of recombination events, we investigated whether this could be explained by difference in the localization of crossover hotspots by comparing that of the HRIs (see above). We first defined “hot windows” as genomic regions that harbour a HRI in at least one population. Figure 5A represents the proportion of the 5,881 resulting hot windows including HRIs that are population specific (HRI in one population only) or shared by two, three or all four populations. Around 66% of these windows are population-specific and 34% are shared by two populations or more. The proportion of hot windows shared by three or four population drops to 12% and 2% respectively. Location of shared HRIs along the genome followed the density of HRIs per region, as most (76%) shared windows were located in telomeric regions R1 and R3 and the rest (24%) in pericentromeric regions R2a and R2b (chisq.test P-Value = 0.06). To check if such an overlap across populations can be explained by chance alone, we compared the observed repartition of hot windows to a simulated distribution obtained by a random assignment of HRIs corresponding to the null hypothesis of the absence of HRI population sharing (see methods). The proportion of common hot windows under this random assignment is represented by grey boxplots on Figure 5A. The observed proportion (coloured points) was always significantly different to the expected proportion under random assignment of HRIs. On average, 95% of hot windows are population-specific if assigned randomly, much more than the 66% we observed. In addition, four-population overlaps were rare in the simulations (8.1% of our simulations) and when they happened they concerned only one or two intervals while we found 139 windows where HRIs are shared between the four landrace populations.

**Figure 5:**
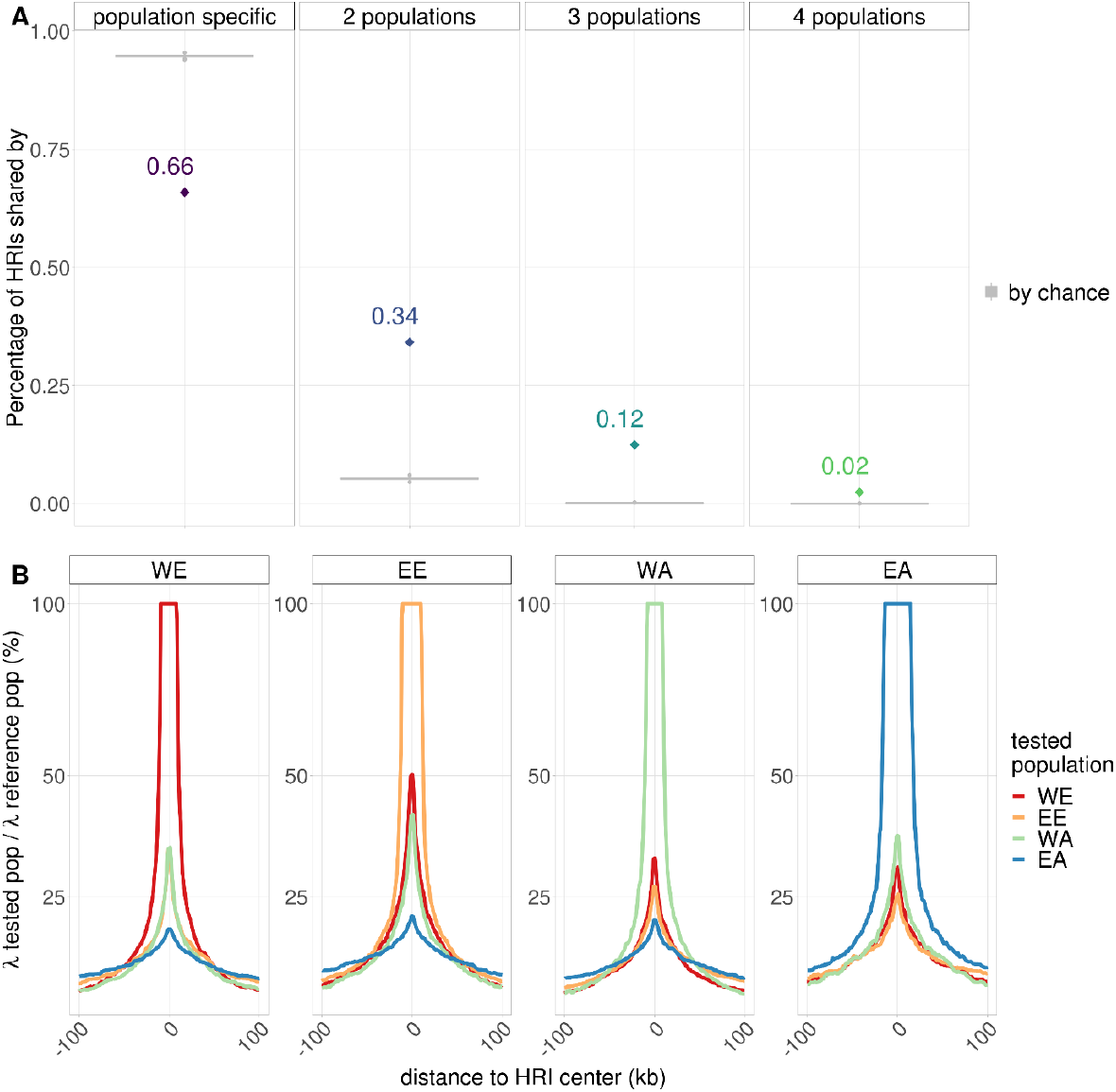
Conservation of highly recombining intervals (HR) across landrace populations. **A** Proportion of co-localizing HR (coloured points) and simulated co-localizing values under random assignment of HRIs (grey boxplots) **B** LD-based recombination intensity in each of the four populations WE, EE, WA and EA around HRIs specific to one population

HRIs shared by more populations tend to be more intense. For example, 55% of WE’s HRIs (λ ≥4) co-localize with HRIs of other populations (λ ≥4), but this proportion rises to 78% when subsampling WE’s HRIs with a higher threshold of λ ≥ 20. The intensity of recombination in a hot window increases when it is shared by more populations: the median of λ is 10.7, 8.1 and 6.9 when shared by 4, 3 and 2 populations respectively and is only 5.9 for population-specific hot windows. This approach to compare HRI between populations depends on the threshold to claim HRIs and our ability to detect them, which can vary between populations. To make up for these effects, we looked at the recombination intensity (λ) observed in one population around HRIs detected in another population (supplementary file S4). Figure 5B presents this average recombination intensity for HRIs detected in each of the four populations. It shows that the local intensity at an HRI position in the other populations is almost twice the background intensity (defined as the intensity measured at 100 kb from the HRI centre (average ratio at HRI positions: 29%; average background ratio: 13%). This further shows that HRIs tend to be shared across populations. Comparing HRI localizations and intensities in all four populations demonstrate that there is a significant amount of sharing of HRIs that could be due to an underlying partial conservation of recombination hotspots.

Further examination of the increase in recombination intensity on Figure 5B reveals that HRI intensities tend to be more similar when populations are more related. For example, around WE’s HRIs, the recombination intensity increases in all populations, but slightly less in EA which is the most genetically distant population to WE. To study this further, we studied quantitatively the relationships between the similarity in recombination profiles and the genetic divergence of populations. To do so, we fitted a linear regression to estimate the effect of the local differentiation index (F_st_) on the similarity of recombination profiles (measured by their correlation) for all genomic regions (R1, R2a, R2b, and R3) on all chromosomes (1A to 7D) (Figure 6). We found that most effects (slopes) were negative, revealing a striking pattern where the similarity in recombination intensity decreases proportionally with genetic divergence: almost all genomic regions (67 among 84) had a negative slope estimate significantly different from 0 and others genomic regions (15 among 84) had negative but non-significant slope estimates different from 0. We stress out here that the similarity in recombination profiles is based on the relative local recombination intensity (parameter λ) that should not be affected by the evolutionary history of populations and that we calculated F_st_ from haplotypes rather than single SNP to avoid an ascertainment effect, although results based on SNP F_st_ showed the same pattern (supplementary figure S8). To further evaluate if the decreasing similarity of recombination patterns could be explained by the varying proportion of shared polymorphisms between population pairs, *i*.*e*. SNP ascertainment, we carried out all our analyses on a subset of 100,381 SNPs that are polymorphic in all four populations. We found that the decreasing similarity of recombination intensities with genetic divergence still hold using this common SNP dataset (supplementary Figure S9), even if the absolute values of slope estimates were smaller (supplementary Figure S10). Finally, these results demonstrate that the similarity in recombination profiles of bread wheat populations is strongly negatively associated with their genetic divergence and highlight that recombination landscapes in bread wheat have been evolving during the establishment of the current genetic structure of wheat populations for reasons that we now discuss.

**Figure 6:**
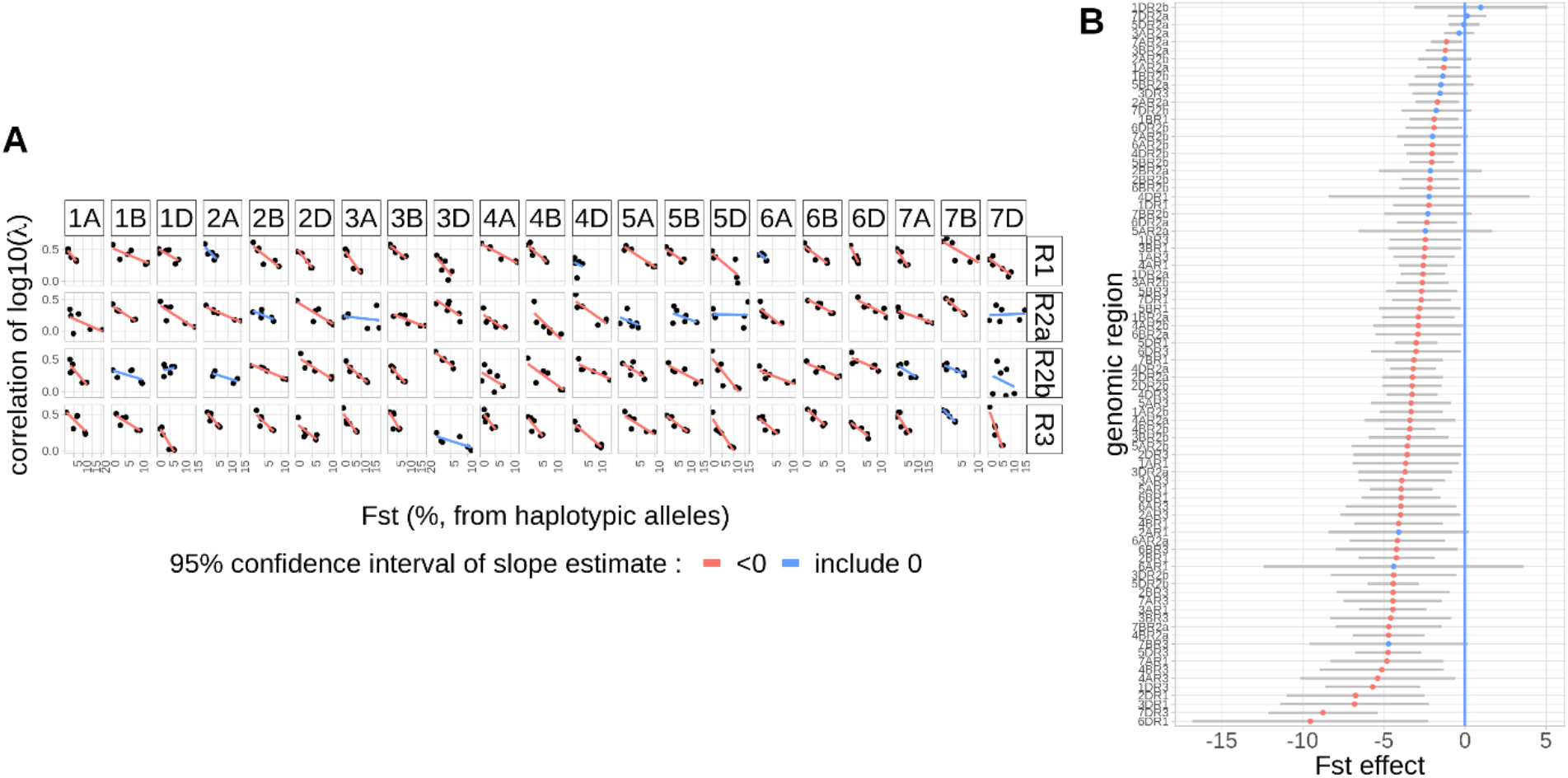
Relationships between correlation of local recombination intensity and F_ST_ per genomic region. **A** Relationship per genomic region. **B** Ranked slope estimates (coloured points) and their 95% confidence interval (grey bar). Blue colour represents slopes with a confidence interval overlapping 0 and red colour confidence interval not overlapping 0.

## Discussion

### LD-based recombination rates can be computed genome-wide in bread wheat

In our study, we estimated LD-based recombination rates for the first time at the whole-genome scale in bread wheat. Previous studies were done at local scale only (Darrier et al. 2017) but suggested that this approach could be applied genome-wide. We used four diverging populations of landraces representative of the four main worldwide genetic groups (Balfourier et al. 2019). For all maps, 80% of the genetic distance was found in 36% (±1%) of the physical distance. This is less concentrated than what was previously observed on single chromosome 3B (80% in less than 20%; Saintenac et al. 2009; Darrier et al. 2017). This discrepancy is likely due, on one hand to the higher SNP density in previous studies on chromosome 3B that allowed to precisely delimit recombination hotspots on this particular chromosome, and on the other hand likely because classical empirical estimates of recombination rates in biparental maps let most of the genome depleted of recombination. However, and as expected, historical crossovers tend to accumulate in distal sub-telomeric regions of the chromosomes (namely R1 and R3 regions). In most organisms, pairing initiation between homologues occurs in many places along the chromosomes but tends to be favoured by a meiosis-specific organisation called “bouquet” where telomeres are gathered on the internal nuclear envelope at the Leptotene stage, just before synapsis (Zickler and Kleckner 2015). The bouquet would then facilitate alignment between homologues and pairing would be simultaneously favoured through the repair of double-strand breaks including crossovers (Zickler and Kleckner 1998; reviewed in Scherthan 2001 or Harper et al. 2004). In bread wheat, distal crossovers would then be predominant because of the bouquet and be limited in R2a and R2b regions because of interference (Saintenac et al. 2009).

At a fine scale, LD-based maps revealed that 1% to 2% of intervals of telomeric and pericentromeric regions (depending on the population) exhibited especially high recombination rate (HRIs), suggesting that these intervals overlapped recombination hotspots. The accumulation of crossovers in recombination hotspots was already observed in bread wheat (Saintenac et al. 2011; Darrier et al. 2017) and seems to be a common phenomenon across many species (for a review see Stapley et al. 2017). Recombination hotspots are usually found to be associated with open-chromatin signatures (for a review, see Dluzewska et al. 2018). In previous study in bread wheat, recombination hotspots were found to locate nearby gene promotors and terminators. Our results are consistent with this finding, as most (80%) of our HRIs are located nearby gene features.

### LD-based recombination maps correlate well with the biparental genetic map

In principle, LD-based recombination maps are suited to study the similarity of recombination profiles of diverging populations. In our study, they allowed to compare recombination rates of four populations with about twice more SNPs than the densest genetic maps currently available (131-170k SNPs in EA and WA respectively, versus 80k SNPs in Rimbert et al. 2018, 55k markers in Liu et al. 2020, 50k SNPs in Jordan et al. 2018). Moreover, LD-based maps are representative of a whole population and less susceptible to individual specific variation, for example introgressions which are known to prevent local formation of CO between the introgressed chromatid and the native chromatid. Introgressions from wild relative species are frequent in bread wheat species, representing from 4 to 32% of bread wheat genome (Zhou et al. 2020).

The limitation of LD-based maps relies on the fact that they can be affected by evolutionary patterns, which in turn can hinder their usefulness to study the evolution of recombination rate. Indeed, to the extent that evolutionary forces and past demographic events (bottlenecks, population expansions, hidden structuration) affect LD patterns they can also affect recombination rate estimates. To measure to which extent LD-based recombination rates differ from meiotic ones, we compared LD-based maps to the CsRe meiotic map. This revealed that, genome-wide, the correlation between the two approaches was very high (≥ 0.7, Table 1). Although part of this correlation is explained by the large differences in recombination rate between chromosomal regions (R1, R2a, R2b, R3 and C), our results also indicate a substantial high correlation within each of these regions. The correlation between LD-based and the CsRe genetic map ranged from 0.50 on average in EA, 0.55 in WA and EE and 0.58 in WE at 4 Mb per genomic region considering all populations but only telomeres and pericentromeres (Table 1). This value is consistent with correlation values obtained in the literature for other plant species. For example, the correlation between LD-based and meiotic recombination map was found to be 0.3 in rice (Marand et al. 2019), 0.81 in barley (Dreissig et al. 2019) and 0.44–0.55 in Arabidopsis (Choi et al. 2013). The correlation values we report are thus likely to be underestimates of the true values. To compute these correlations, we used estimates of recombination rates. Like any statistical estimates they come with measurement errors of the true parameters. Hence the correlation between estimates, providing these errors are independent, are necessarily smaller than the true correlation (Fisher 1915). Apart from this statistical effect, we could also explain some of the differences between LD-based maps and the meiotic map by genomic rearrangements (introgressions on chromosome 7D and 2A in Renan) that are specific to the CsRe population: in these regions the CsRe recombination profile is not representative of the landraces recombination profiles.

The overall similarity between the meiotic map and LD-based maps shows that LD-based recombination patterns offer a robust representation of the distribution of recombination along the bread wheat genome.

### Robustness of LD-based recombination maps

Despite good concordance with the meiotic map, LD-based recombination maps can still be locally affected by demographic effects, and thus result in bias when interpreting differences or similarities between populations. For example, Kim and Nielsen (2004) and Chan et al. (2012) showed that selective hard-sweeps can produce LD patterns that mimic those of recombination hotspots. Dapper and Payseur (2018) showed that demographic events can decrease the power to detect hotspots leading to an under estimation of the co-localization of LD-based recombination hotspots when using LDhat software (Auton and McVean 2007). Here, we used PHASE (Li and Stephens 2003), a software to infer recombination rates from LD patterns that implements a quite different methodological approach than LDhat but it is possible that its inference is also affected by such effects. In particular, there were twice many HRIs detected in WE (2,739) and WA (2,743) than in EE (1,968) and EA (1,253), representing a significant variation from 1% of intervals in EA (122,490 SNPs once centromeres removed) to 2% of intervals in WE (161,953 SNPs once centromeres removed) (significant chisq test, P-value < 2.2e-16). Although this varying number of HRIs per population could result from a variation in recombination patterns, it is likely also due to differences in the power to detect HRIs in each population which would be consistent with results from Dapper and Payseur (2018). Indeed, as the proportion of HRIs per population follows the levels of admixture and SNP density (both higher for WE and WA than for EE and EA), this favours a possible contribution of a different detection power to the variation of HRIs per population. However, we did not observe any atypical LD-based estimate for intervals located nearby genes known to be involved in domestication (*e*.*g*. brittle rachis, tenacious glume, homoeologous pairing or non-free-threshing character) or in recent crop improvement. To further reduce the potential influence of demographic forces on our inference, we performed the comparison between population maps, not on LD-based recombination rates themselves (ρ) but on the relative rate (λ) of recombination in an interval compared to its neighbours in windows of 2 centi-Morgans. Using relative rates should clean our inference from any local effect of demographic forces, especially selection that could tend to be more shared between closely related populations than distant ones.

Results were not much affected by SNP ascertainment or the method used to calculate the F_ST_ index. The decreasing similarity of recombination rates with genetic differentiation still hold when estimating LD-based recombination rates on a population specific SNP dataset or a common SNP dataset. The co-localization of highly recombining intervals was also not influenced by the SNP dataset (supplementary Figure S11). The estimation of F_ST_ index, using either haplotypic or SNP alleles, provided also consistent results. Overall, these results strongly support the idea that the decrease of similarity in LD-based recombination profiles is not an artefact of demographic forces or biases due to SNP ascertainment but that the underlying recombination profile is linked to the divergence of populations.

### Evolution of the recombination landscape in bread wheat

Our results are consistent with previous reports. Gardiner et al. (2019) showed that closely-related bread wheat parental lines lead to RIL populations with more similar crossover profiles. (Darrier et al. (2017) compared LD-based recombination profiles of a European and an Asian population, the two main ancestral bread wheat genetic pools, on two scaffolds of 1.2 and 2.5 Mb on chromosome 3B. They found that LD-based recombination profiles are broadly conserved, but highlighted that hot intervals in LD-based recombination profiles were not necessarily shared between these two pools. Similar results were observed in other plant species such as rice (*Oryza sativa*; Marand et al. 2019) and cocoa-tree (*Theobroma cacao*; Schwarzkopf et al. 2020). Other plant studies hint at a possible decreasing similarity of fine scale recombination profiles over evolutionary time measured by F_ST_ (maize *Zea mais* Rodgers-Melnick et al. 2015); poplar *Populus* species Wang et al. 2014; Wang et al. 2016); cotton *Gosypium hirsutum* Shen et al. 2019); barley *Hordeum vulgare* Dreissig et al. 2019).

Several hypotheses can be formulated to explain the differences in recombination profiles between populations. First, this can be due to environmental effects. This is the case in barley, where recombination rates vary along the genome and are affected by environmental conditions as well as by domestication (Dreissig et al. 2019). For example, high temperatures are known to affect meiosis and above 35°C, this may lead to complete failure and severe sterility (Loidl 1989; Higgins et al. 2012). Interestingly, within a range of 22-30°C, highest temperatures may modify the recombination profile. In barley, it was shown that at 30°C, distal recombination events are reduced while interstitial events became more frequent revealing thus a slight shift and a modification of the global recombination profile (Higgins et al. 2012). However, in our case, this hypothesis is not the most likely as we were using populations from the same hemisphere and latitudes, with landraces from different countries. Environment is thus certainly very different between all the origins of our landraces and temperature should vary a lot in each location and is not stable enough to affect durably and maintain a different recombination profile between the four populations. Moreover, it was recently shown that increased temperature up to 28°C for three weeks during wheat meiosis has only a limited impact on recombination distribution (Coulton et al. 2020).

Secondly, differences in recombination profiles can be explained by differences in the chromatin accessibility landscape during meiosis between populations. Many studies showed that chromatin status is the main feature that drives recombination in plants. DNA is partitioned in blocks of heterochromatin and euchromatin which are dispersed along the chromosomes. In bread wheat, heterochromatin preferentially locates in pericentromeric regions while euchromatin-rich DNA is more frequent in distal subtelomeric regions of the chromosomes (IWGSC 2018). In Arabidopsis, it was shown that crossovers are enriched in euchromatin and mainly occur close to genes promoters and terminators (Choi et al. 2013; Drouaud et al. 2013). Meiotic recombination profile in this species is also shaped by H2A.Z nucleosome occupancy, DNA methylation or epigenetic marks such as Histone 3 Lysine 9 di-methylation (H3K9me2; (Choi et al. 2013; Underwood et al. 2018). This led to our second hypothesis that chromatin status has evolved between our four populations, rather than an evolution of the recombination determinism itself. It is likely that during the evolution process, there has been a selection pressure around different genomic regions depending on geographical area. This selection pressure could therefore contribute to the deposition of histone landmarks to regulate gene activity such as H3K4me3, H3K9ac and H3K27ac that are associated with transcriptional activation (Roth et al. 2001; Howe et al. 2017) or on the contrary H3K27me3 and H3K9me3 associated with transcriptional suppression (Saksouk et al. 2015). Interestingly, in some mammals, recombination is directed by the zinc finger protein PRDM9 that possesses a set domain that catalyses the trimethylation of lysine 4 of H3 to produce H3K4me3 (for review see Grey et al. 2017). Similar mechanisms involving histone 3 modifications such as methylation or acetylation that could affect recombination profile afterward are thus likely in plants.

Another factor that may explain the difference of recombination patterns between the populations could be the natural introgression of alien DNA fragments from wheat relatives during the evolution process. Introgressions from wild-species have been widely used and more than 50 alien germplasms have been used to improve wheat varieties (Wulff and Moscou 2014). For example, Renan possesses two introgressed fragments from *Aegilops ventricosa* conferring resistance to leaf, yellow and stem rusts (*Lr37*/*Yr17*/*Sr38*) on chromosome 2A (2A/2N translocation) and to eye-spot (*Pch1*) on chromosome 7D (7D/7Dv translocation; Maia 1967; Helguera et al. 2003). These introgressions repress recombination (Worland et al. 1988) and this resulted in a poor correlation between CsRe genetic map and our LD-based maps for genomic region 7DR3 in our analysis. It was recently shown that natural or artificial introgressions of wheat wild-relatives DNA contributed to up to 710 Mb and 1580 Mb in wheat landraces and varieties respectively (Cheng et al. 2019), and represent from 4 to 32% of bread wheat varieties genome (Zhou et al. 2020). A similar analysis used exome capture to evaluate introgression in 890 hexaploid and tetraploid wheats (He et al. 2019). The results also suggest that introgressions of DNA fragments from wheat relatives contributed significantly to improve the diversity of current wheat cultivars. Since natural introgressions are frequent in wheat landraces and because they contribute to modify the recombination profile, we could hypothesise that these introgressions are different in our four collections, which would result in different recombination profiles as well. Only an extensive sequencing of our accessions would allow to bring the answer.

## Conclusion

This study demonstrates the evolution of the recombination profile at a genome-wide scale in closely-related wheat populations with increasing genetic divergence. Based on recombination landscapes robust to demographic events, the comparison of the four landrace populations revealed a clear signal of a decreasing similarity between fine-scale recombination landscapes with increasing genetic divergence. Specifically, we found (i) that highly recombining intervals were more shared between closely related populations, (ii) recombination intensities at HRIs detected in one population decreased in the other populations with their genetic divergence and (iii) the correlation of recombination landscapes between pairs of population decreases with their local genetic distance as measured by F_ST_. Our results, interpreted in the light of previous findings in bread wheat and other species, clearly shows that recombination landscapes in wheat change with genetic divergence between populations. Being based on closely related populations that recently diverged (no more than 10 000 years ago), this study further shows that this divergence can be quite fast. Reasons for this divergence remain to be found but our results can hint at some possibilities. Further analyses are needed to settle this question, which should greatly help developing original approaches useful for wheat improvement and breeding.

## Materials & Methods

### Plant material

A collection of 632 bread wheat landraces (Balfourier et al. 2019) was genotyped on the TaBW410k SNPs, including 280k SNPs from the Axiom Affymetrix® TaBW280k SNP array. Besides, a population of 406 F6 Recombinant Inbred Lines (RILs) derived from the cross between the Asian variety Chinese Spring and the European variety Renan (CsRe), were also genotyped on the TaBW280k SNP array. After quality filtering including control of missing data rate (10% maximum), heterozygosity rate (5% maximum), excluding off-target variants (OTVs), 578 landraces genotyped with 200,062 SNPs were kept for the population-based analysis and 79,564 polymorphic SNPs were successfully mapped on the CsRe population.

The physical positions of SNPs on the 21 bread wheat chromosomes were determined using BLAST (Basic Local Alignment Search Tool; Altschul et al. 1990) of context sequences on the International Wheat Genome Sequencing Consortium RefSeq v1.0 genome assembly (IWGSC 2018). Position of high confidence genes, exon, 5’UTR and 3’UTR were extracted from RefSeq V1.0 annotation.

### Robust estimation of the meiotic recombination profile

Due to the relatively low number of meiosis sampled in the CsRe data, a Bayesian model inspired from Petit et al. 2017 was used to obtain robust estimates of recombination rates. We modelled the probability distribution of the recombination rates observed in RIL (*C*_*i*_) given the number of observed recombination events (*y*_*i*_) as:

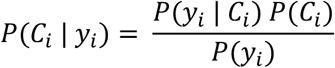

The likelihood *P*(*y*_*i*_|*C*_*i*_)is modelled as a Poisson distribution, its parameter being the expected number of recombination events in an interval and computed as: 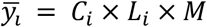, where *L*_*i*_ is the physical size (in megabases, Mb) of the interval and M the total number of RILs. Thus, the likelihood of the recombination rate *C*_*i*_ is:

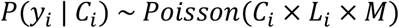

To specify a prior distribution of *P*(*C*_*i*_), we considered that the wheat recombination landscape varies widely along a chromosome. According to the nomenclature of Choulet et al. 2014, each of the wheat chromosomes can be segmented into five chromosomic regions associated with different global recombination rates and genomic content: two highly-recombining telomeric regions (R1 and R3), two low-recombining pericentromeric regions (R2a and R2b) and one centromeric region (C) where recombination is almost completely suppressed. The small arm of each chromosome is composed of R1 and R2a while the long arm is composed of R2b and R3. The physical size of these regions ranges between 10 Mb for the smallest telomere to 321 Mb for the largest pericentromere (supplementary file S5). To account for the specific range of recombination rate variation in each region in our model, the prior distribution of the recombination rates in each of these regions was a specific Gamma distribution:

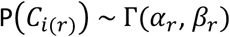

where r denotes the region,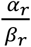 gives the mean of the Gamma distribution and 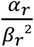 gives the variance. The Gamma distribution being a conjugate prior to the Poisson distribution, the posterior distribution of *C*_*i*_ is also a Gamma distribution:

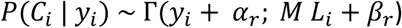

The posterior mean of *C*_*i*_ (in M/Mb) is then:

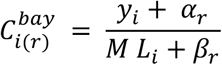

The parameters *α*_*r*_ and *β*_*r*_ of the prior Gamma distribution were set using an empirical Bayes approach, (*i*.*e*. estimating prior distribution directly from data), independently for each of the five r regions (supplementary Figure S12). A Gamma distribution was fitted (R MASS package, Venables and Ripley 2002) over the distribution of empirical recombination rates observed in RIL. This latter was computed as

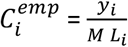

Note that null recombination rates were replaced by the lowest non-null estimate of recombination rates of the region to allow fitting the Gamma distribution. We derived the meiotic recombination rates from the RILs recombination rates using the Haldane and Waddington formula (Haldane and Waddington 1931) and the Morgan mapping function (cM = frequency of recombinants * 100). Indeed, the size of intervals (median = 5 Kb) were small enough to consider that interference is very strong within and thus one recombination in one individual result from only one crossover (and not from coincidence of several crossovers). We thus obtained the Bayesian meiotic recombination rate 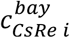 (cM/Mb).

#### Considering uncertainty in crossover locations

For estimation of recombination rates, it was necessary to count the number of recombinants in CsRe intervals (*y*_*i*_). Missing data on genomic segments with no parental allele switch at segment extremities were imputed. A crossover was counted at each parental allele switch, yielding 26,239 crossovers. Due to the presence of missing data in RIL genotypes, a number of switches did not occur between pairs of immediately adjacent markers. In such cases, the crossover cannot be assigned with certainty to a single interval of two successive SNPs. For example, a RIL genotype AA/--/BB identifies a switch between the first and third marker but cannot discriminate a recombination in the first vs. the second interval. In such cases, we accounted for the uncertainty in crossover location following the sampling procedure of Petit et al. (2017). Briefly, each crossover is overlapped by a set of one or more intervals. A sampling procedure assigned each crossover to a particular interval with a probability computed as the size of the interval divided by the size of the crossover area (physical distance between the two closest SNPs showing different parental alleles). Repeating 1,000 times the sampling procedure yields 1,000 estimates of *y*_*i*_ per interval, which can then be converted into recombination rates and averaged.

### LD-based recombination profiles of four diverging populations of landraces from patterns of linkage disequilibrium

#### Identification of diverging populations of landraces representative of bread wheat worldwide diversity

Balfourier et al. (2019) analysed the genetic structure of the landrace dataset and could pinpoint four main groups corresponding to the geographic origins of lines. Despite this structuration, the general pattern of differentiation in these data is somewhat continuous, a lot of individuals exhibiting admixed origins. Here, we subsampled the dataset in order to constitute populations of individuals that were both homogeneous within groups and clearly differentiated between groups. This was done in two steps that we now describe.

From the Balfourier et al. 2019 admixture analysis with K=4 groups, landraces exhibiting an admixture coefficient smaller than 50% of their dominant group were removed, yielding 534 “low admixed” landraces. These 534 landraces were grouped (again) into 4 populations by hierarchical clustering on the pairwise distance matrix estimated in Balfourier et al. (2019) and using the Ward’s grouping criterion. The four populations were named as West Europe (WE), East Europe (EE), West Asia (WA) and East Asia (EA) from the geographical origin of their members. The genetic distance between two landraces was the proportion of mismatched haplotypic alleles along the genome, computed using 8,741 haplotypic blocks containing up to 20 alleles per block (Figure 1 of Balfourier et al. (2019)). After this first step, the second one was aimed at discarding closely related individuals within each population to avoid over representing family specific recombination events. Pairs of individuals exhibiting a very low genetic difference were discarded, keeping 371 landraces (more details in supplementary protocol and supplementary Figure S13).

#### Evolutionary distance between populations measured by F_ST_

Pairwise differentiation indexes (F_ST_) of the four populations were computed within each genomic region (chromosomal region within a chromosome *e*.*g*. 1AR1) using alleles of 8,741 haplotypic blocks (Weir & Cockerham distance, R hierfstat package, function pairwise.WCfst, Goudet and Jombart 2015) or SNPs (Reynolds distance, HAPFLK software, Bonhomme et al. 2010; Fariello et al. 2013) (supplementary file 6).

#### Inferences of LD-based recombination rates from linkage disequilibrium patterns

LD-based recombination rates were estimated using PHASE software V2.1.1 (Li and Stephens 2003; Stephens and Scheet 2005). PHASE inputs were successive windows of SNPs along the genome, constituted of one central part and two flanking parts overlapping the previous and the next windows to avoid border effect in PHASE inferences. Central and flanking parts spanned on average 1 cM and 0.5 cM respectively based on the CsRe genetic map (supplementary protocol S3). PHASE was run for each window with default options, except for two parameters of the Markov Chain Monte Carlo (MCMC), following recommendations of the documentation on estimating recombination rates. The number of sampling iterations was increased to obtain larger posterior samples (option -X10) and the algorithm was run 10 times independently (option –x10) to better explore combinations of parameters and keep the run with the best goodness of fit. The sampling stage of the MCMC yielded 1,000 samples of the posterior distribution of:

- The background recombination rate of the window w: *ρ*_*w*_
- The ratio *λ*_*i*_ between the background recombination rate of the window *ρ*_*w*_ and the LD-based recombination rate in each interval i of two successive SNPs *ρ*_*i*_ so that *ρ*_*i*_ = *λ*_*i*_ * *ρ*_*w*(*i*)_where *w(i)* identifies the window which interval *i* belongs to. The parameter *λ*_*i*_ can be seen as a measure of local recombination intensity compared to genomic background (inflation or deflation).

PHASE samples jointly *ρ*_*w*_ and *λ*_*i*_ in their posterior distribution at each iteration, so their product yields 1,000 samples of the posterior distribution of LD-based recombination rate *ρ*_*i*_ (/kb) (supplementary file S7).

#### Correlation of LD-based recombination profiles

To compare LD-based recombination profiles, it was necessary to obtain a common set of intervals across the four populations (WE, EE, WA, EA), as polymorphic SNP sets were different. We defined smaller intervals formed of successive markers that were polymorphic in at least one population (supplementary Figure S14). For each population, the recombination estimates in smaller intervals were considered to be the same as the estimates belonging to population specific intervals overlapping them, assuming that recombination rates are constant within intervals. We removed intervals not overlapped by all populations on chromosome extremities. This process yielded a complete factorial dataset of 194,409 intervals with no missing data and a set of 1,000 values sampled from the posterior distribution for each parameter *ρ*_*pi*_ and *λ*_*pi*_ per interval *i* and per population *p* (supplementary file S8). The similarity between LD-based recombination profiles was measured by correlating the *log*_10_ of median of *λ*_*pi*_ (noted 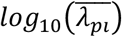) of all intervals between different populations. The median of posterior distribution of *λ*_*pi*_ was chosen as a it is robust to outliers in the posterior distribution, as recommended (Li & Stephens 2003) and using the log scale is natural when comparing intensities across groups. To obtain correlation coefficients, a linear model including a full unstructured variance-covariance matrix was fitted on 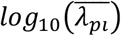, so that each population had its own range of variation of local recombination intensity and each pair of population has a specific covariance parameter:

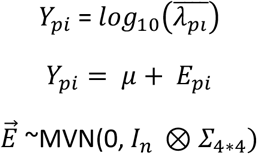

where *∑*_4*4_ is a variance-covariance matrix from which we extract correlation coefficients.

The model was applied independently to each genomic region (from 1AR1 to 7DR3, except centromeric regions). The total number of intervals n per genomic region ranged from 154 to 8,131. The differences of recombination intensity profiles across the four populations of landraces was assessed by model comparison. We compared the Bayesian Information Criterion (BIC) of a model with a full variance-covariance matrix to a simpler model with a variance-covariance matrix including only one correlation parameter for all pairs of populations. The complex model was deemed to be a better model if its BIC was inferior to the BIC of the simpler model. The models were fitted with ASREML-R V3 (Butler et al. 2009)

#### Co-localisation of highly recombining intervals between populations

Intervals with a LD-based recombination rate exceeding four-times or more the background recombination rate (λ ≥ 4) figuring as outliers in λ distribution (supplementary Figure S15 and supplementary file S9), were defined as highly recombining intervals (HRIs) and adjacent HRIs within a population were merged. Due to strong heterogeneity of HRI’s size, we discarded too small or too wide HRIs (supplementary protocol S4). For each HRI in each population, the overlapping HRIs in other populations were recorded. A set of HRIs intervals was considered as co-localizing in two, three or four populations if all HRIs overlapped each other (*i*.*e*. they formed a clique in network terminology). Note that this implies that a wide HRI can potentially be involved in more than one clique. For each group of co-localizing HRIs (each clique), we defined a hot window as the smallest common overlapped area (supplementary file S10). Population specific HRIs, *i*.*e*. HRIs which did not overlap any other HRIs, also formed hot windows whose frontiers were defined by the upper and lower limit of HRIs. Each hot window thus included HRIs of one, two, three or four populations. The proportion of HRIs shared by two populations or more (for example WE and EE) was computed as the number of hot windows including HRIs of each population (hot windows including both WE’s HR and EE’s HR) divided by total number of hot windows (including either WE, EE, WA or EA’s HRIs) (supplementary Figure S16). Dividing by the total number of hot windows is more convenient to compare the proportion of HRIs population-specific, or shared by two, three or four populations.

To test for the hypothesis that the observed proportion of HRIs shared by populations is due to chance, an empirical range of plausible values of co-localization due to chance was estimated by simulation. In 1,000 simulations, each HRI of each population was assigned to a random interval within the genomic region it belongs (1AR1 to 7DR3) and the proportion of shared hot windows was computed (supplementary file S11).

### Comparison of the LD-based recombination rates and the CsRe meiotic recombination rates

The comparison between meiotic (CsRe) and LD-based recombination rates were done on windows of 4 Mb (∼ 1 cM on average, wide enough to accurately estimate intrinsic recombination rate) along the genome. Meiotic recombination rates were estimated using the Bayesian model described above, the attribution of crossover to windows being done using the (Petit et al. 2017) approach (see above). To compute the LD-based recombination rate in 4 Mb windows, the total LD-based genetic distance per window of 4 Mb was divided by the total physical distance and averaged over the 1,000 samples of the posterior distribution:

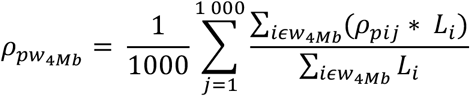

with *i* the interval and *j* one posterior distribution value among 1,000 (supplementary file S12).

## Acknowledgements

The authors would like to thank to the genotoul bioinformatics platform Toulouse Occitanie (Bioinfo Genotoul, doi: 10.15454/1.5572369328961167E12) for providing help, computing and storage resources. The authors thank Hélène Rimbert for her help in identifying SNP positions on refSeq v1.0 and with the CsRe and refSeq v1.0 data files, Ingrid David for her help with the ASREML software and Gilles Charmet and Jean-Michel Elsen for their advices on the manuscript. Doctoral work of ADDD was funded by the INRAE metaprogram SELGEN and Florimond Desprez (Cappelle-en-Pévèle, France). This work was supported by the Bread Wheat grant (ANR-10-BTBR-03).

